# Malaria during pregnancy and newborns outcome in an unstable transmission area in Brazil: a population-based record linkage study

**DOI:** 10.1101/244178

**Authors:** Jamille Gregório Dombrowski, Rodrigo Medeiros de Souza, Natércia Regina Mendes Silva, André Barateiro, Sabrina Epiphanio, Lígia Antunes Gonçalves, Cláudio Romero Farias Marinho

## Abstract

**Background:** Malaria during pregnancy is one of the major causes of mortality in tropical regions, causing maternal anemia, intrauterine growth retardation, preterm birth, and low birth weight (LBW). The integration of the information systems is crucial to assess the dimension of gestational malaria in a wide and useful way, to improve decision making and maternal-child health.

**Methods and Findings:** An observational population-based study acquired information retrospectively from all live births that occurred between 2006 and 2014 in Cruzeiro do Sul (Acre, Brazil). Social and clinical data of the mother and newborn was extracted from the Information System of Live Births. Malaria episodes information was obtained from the Brazilian Epidemiological Surveillance Information System Malaria. A deterministic record linkage was performed to assess malaria impact on pregnancy. The studied population presented a malaria incidence of 8.9%, of which 63.9% infected by *Plasmodium (P.) vivax.* Reduction of newborns birth weight at term (small for gestational age (SGA) and LBW) has been found associated with *P. vivax* infection during pregnancy (SGA - OR 1.24, 95% CI 1.02-1.52, p=0.035; term LBW - OR 1.39, 95% CI 1.03-1.88, p=0.033). Additionally, *P. falciparum* infection during pregnancy has been found to be associated with preterm births (OR 1.54, 95% CI 1.09-2.18, p=0.016), which is related with late preterm births (OR 1.59, 95% CI 1.11-2.27, p=0.011).

**Conclusions:** Despite the decrease of malaria cases during the evaluated period, we present evidence of the deleterious effects of gestational malaria in a low transmission area in the Amazonian region. Regardless of *Plasmodium* species, malaria during pregnancy poses a risk for newborns birth weight reduction, highlighting the impact that *P. vivax* has on the fetus.

**Funding:** São Paulo Research Foundation - FAPESP/Brazil.

## INTRODUCTION

Malaria is a severe and potentially fatal parasitic disease that constitutes a major public health issue, being one of the greatest causes of mortality in tropical regions. Pregnant women are particularly vulnerable to malaria infection and are estimated that 125 million women are at risk of malaria in pregnancy each year ^1^. Malaria can be devastating for both mother and fetus, leading up to 10,000 maternal and 75,000 to 200,000 child deaths each year ^2^. Maternal malaria presents a significant impact on the neonates, being the primary cause of abortion, stillbirth, premature delivery, fetal death, low birth weight (LBW) and fetal/child development retardation in malaria-endemic countries ^2^.

LBW reflects an intra-uterine growth retardation (IUGR) and preterm delivery, which are compelling indicators of infant morbidity ^2-5^. LBW has been linked to infant mortality and poor cognitive development, and the occurrence of non-communicable diseases later in life ^5,6^. In fact, LBW in newborns due to malaria is related with up to 100,000 infant deaths each year in endemic countries ^7,8^. These adverse birth outcomes have been extensively associated with *P. falciparum* infection during pregnancy. In contrast to *P. falciparum,* the *P. vivax* burden in pregnancy is less well described, and have been described as having less impact in the newborn ^2,9^. Though, recent studies have presented the two species as similar threats to the mother and fetus ^10^. Despite the efforts to reduce malaria the prevalence of these adverse birth outcomes remains high.

Therefore, it is crucial to have an efficient epidemiological surveillance of malaria during pregnancy. The linkage of two or more health public surveillance record databases with shared variables presents an important and effective strategy to plan preventive measures. Currently, most of the malaria-endemic countries have malaria public surveillance record databases since it is compulsory notification disease. This will contribute to the identification of epidemics and areas most affected. Thus, allowing to direct and intensify the control and preventive measures to the affected communities, and reduce negative birth outcomes ^11^. In fact, due to the potential assemble with other information systems it can be recognized as an important tool for research ^12,13,14^.

In 2003, the Brazilian Epidemiological Surveillance Information System (SIVEP)-Malaria was implemented to systematize the flow and quality of the information on malaria. This system gathers information on malaria morbidity according to gender, age, *Plasmodium* species, site of residence, the probable site of infection, treatment, and pregnancy status ^15^. Another essential Brazilian information system of national coverage is the Information System of Live Births (SINASC), implemented in 1990. This system collects and systematizes information on maternal, pregnancy, delivery and newborns data ^16,17^. Linkage of record databases is still scarcely used in Brazil, despite being an easy to perform technique with low operational cost. Here we present the first study that evaluates the association between gestational malaria and adverse birth outcomes in the Brazilian Amazonian region for nine years (2006-2014), using information obtained through the linkage of SINASC and SIVEP-Malaria.

## METHODS

### Study design and data collection

This is a population-based observational study developed in the city of Cruzeiro do Sul - Acre (Brazil), located in the Brazilian Amazonian region (7°37’51’’S, 72°40’12’’W) (Fig 1). Cruzeiro do Sul has an estimated population of 82,075 inhabitants and an average of 1,650 births per year ^18^. Together with Porto Velho and Manaus, the three cities are responsible for 21.9% of the malaria cases notified in Brazil ^19^. It is a high transmission risk city, with an annual parasitic incidence of 214 cases per 1,000 inhabitants, with the prevalence of *P. vivax* infection ^19^. The universe of the studied population was composed of all live newborns delivered by women living in the city, between January 2006 and December 2014. The information regarding the mother, newborn and delivery was extracted from SINASC, and the information on the malaria episodes and parasite species was obtained from the SIVEP-Malaria. By knowing that the primary health care is provided free of cost in Brazil and that the information systems of the Ministry of Health (MoH) offer wide coverage, we can presume that these datasets are reliable.

**Figure 1.**
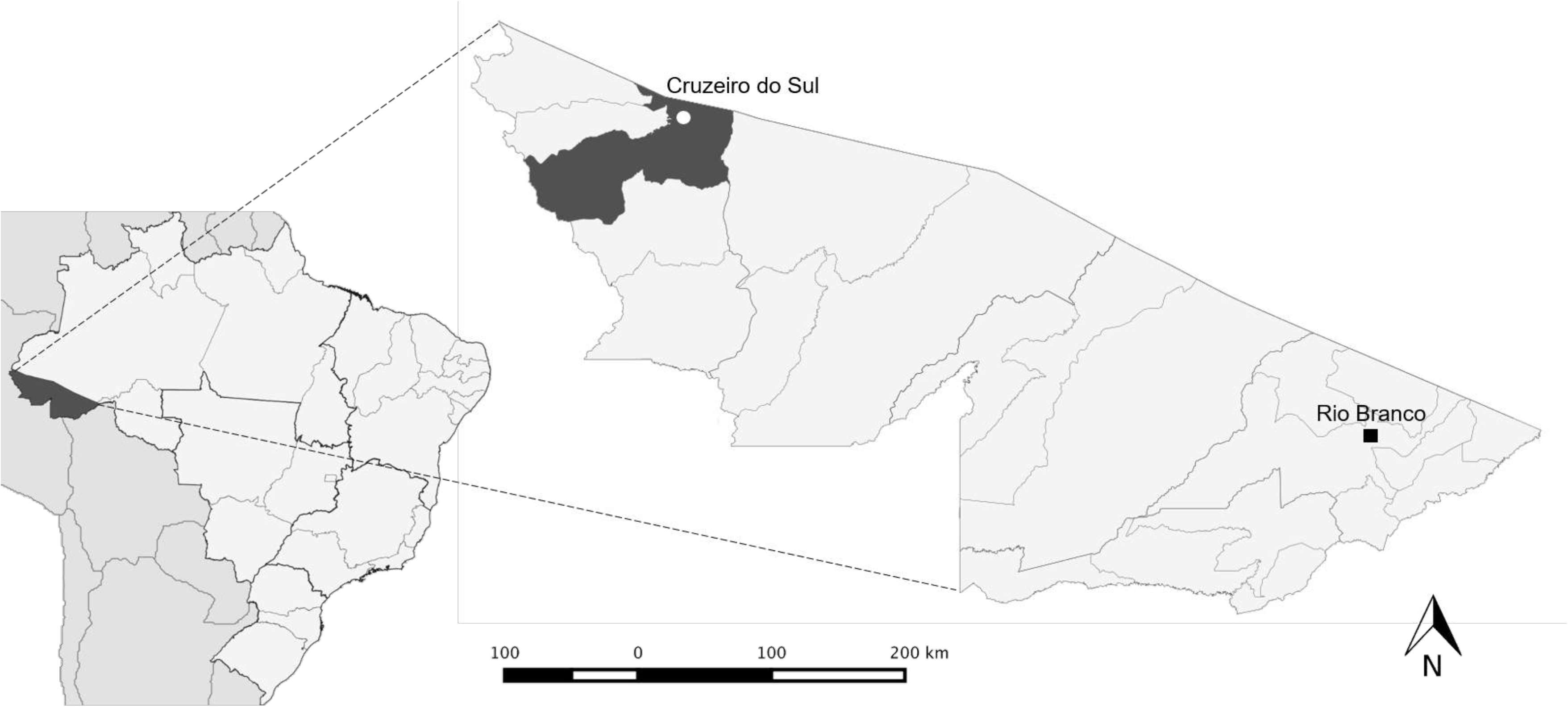
Map showing the geographic location of Cruzeiro do Sul, Acre State, Brazilian Amazon. Cruzeiro do Sul has an estimated population of 82,075 inhabitants. The map also indicates Rio Branco, the capital of Acre state.

### Ethical considerations

According to the Resolution nº 196/96 of the Brazilian National Health Committee, ethical clearance was provided by the committees for research of the University of São Paulo and the Federal University of Acre (Plataforma Brasil, CAAE: 03930812.8.0000.5467 and 03930812.8.3001.5010, respectively). The authors have agreed to maintain the confidentiality of the data collected from the medical records and databases, by signing the Term of Commitment for the Use of Data from Medical Records.

### Exclusion criteria

In this study, the SINASC database was considered the reference. Before performing the record linkage, curation was performed, and newborns with double entries, lack of information on birth weight, presenting congenital diseases or twins were excluded. Upon the linkage of SINASC with SIVEP-Malaria database, newborns with less than 22 weeks (miscarriage) or with no information on the gestational age at birth were excluded (Fig 2).

**Figure 2.**
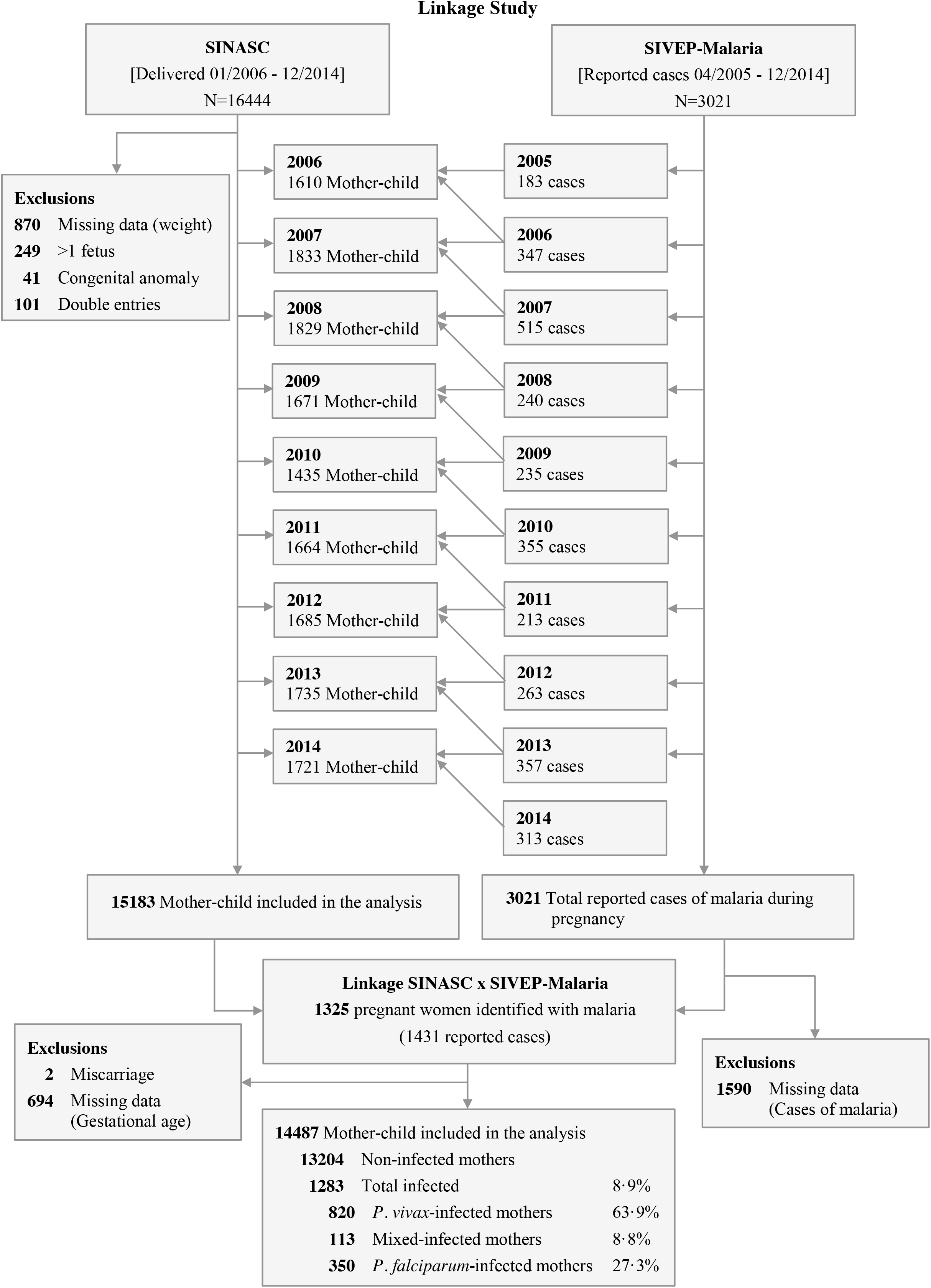
Flowchart detailing exclusion criteria applied to the evaluation of the enrolled maternal-child pairs. Mixed infection - *P. vivax-* and *P. falciparum-infection* occurring at the same time and/or at different times during pregnancy.

### Screening of malaria infection

In Brazil, whenever individual show suspicious malaria symptoms, it is tested by qualified endemic agents that monitor micro-regions. The gold standard method for malaria diagnosis is the thin and thick blood smear, which is screened by trained microscopists from the System of Epidemiological Surveillance of the MoH, and further revised by senior experts, to confirm the results. Infections were categorized per parasite species: *P. falciparum, P. vivax,* or mixed infections. All women who had malaria during pregnancy were treated with antimalarial drugs under medical prescription, according to the Brazilian MoH guidelines.

### Definitions and gestational age estimation

LBW was defined as birth weight < 2500 grams (g). WHO child growth standards were used to classify the small for gestational age newborns, weight ≤ 10^th^ centile (boys ≤ 2758 g, girls ≤ 2678g). The very preterm birth was defined as birth between ≥28 and <32 weeks’ gestation; late preterm birth was defined as birth between ≥32 and <37 weeks’ gestation, and total preterm birth was defined as birth <37 weeks’ gestation. The gestational age was established by the woman’s last menstrual period and, when possible, adjusted by ultrasound during antenatal visit care. In SINASC database, gestational age is categorized as follows: less than 22 weeks’ gestation, 22-27 weeks, 28-31 weeks, 32-36 weeks, 37-41 weeks, and 42 weeks or more.

### Record linkage strategy

The record linkage was performed by using the RecLink III software through the deterministic method (manual search). For the data preprocessing, standardization of both databases was performed by withdrawing accentuations, extra spaces, special characters, and prepositions. After, databases were unified only by two shared variables that presented the appropriate fulfillment. Each year of the record linkage corresponds to one year of the SINASC (containing births records) assembled with two years of the SIVEP-Malaria, to identify all malaria cases presented by women during pregnancy. The linked database gathered the variables from SINASC (mother age, gestational age at delivery, parity, number of antenatal visits, birth weight, and type of birth), with variables from SIVEP-Malaria (infection by *Plasmodium* spp. (yes / no) and parasite species).

### Statistical analysis

Data were extracted into Microsoft Excel, and Stata 14.2 and GraphPad Prism software were used for statistical analyses. We used descriptive statistics to assess the distribution of all continuous (means and standard deviation [SD] or median and interquartile ranges [IQR]), and categorical (frequencies and percentages) variables. Differences between groups were evaluated using Mann-Whitney U-tests, accordingly. Categorical data and proportions were analyzed using chi-square tests. Every p values were 2-sided at a significance level of <0.05. To assess the association between malaria and birth weight reduction or prematurity, adjusted odds ratios (OR) with 95% confidence intervals (CI) were estimated using a multivariate logistic regression approach. These models included infections by malaria (yes / no), maternal age (≥ 18 years old / ≤ 17 years old), gravidity (primigravida / multigravida), and years of formal education (≥ 4 years / ≤ 3 years) as explanatory variables, and birth weight [≤ 10^th^ centile] (yes / no) or LBW (yes / no) as response variables. The first category for each explanatory variable was considered as reference ^20^.

## RESULTS

### Study Population and Baseline Characteristics

Between January 2006 and December 2014, 16,444 births occurred in Cruzeiro do Sul (Acre) with a total of 3,021 malaria cases notified during pregnancy. After applying the exclusion criteria, 14,487 maternal-child pairs remained for further analysis (Fig 2). Table 1 shows maternal characteristics according to infection status (detailed by year in the S1 Table). To highlight that: circa 35% of women were primigravida; above 40% had at least 8 years of formal education (despite the high proportion of no-schooling women); and more than 70% had a minimum of four antenatal visits (Table 1). Nevertheless, it was possible to observe that there were no major differences between non-infected and infected mothers. Malaria incidence in the studied population was 8.9%, with *P. vivax* contributing to 63.9% of the cases (Fig 2). Time series of malaria cases in pregnant women allowed to detect three epidemic peaks along the studied period, one in 2007 with more than 500 cases, and other two in 2010 and 2013 (Fig 3A and S2 Table). Interestingly, the significant reduction of cases from 2007 to 2008 coincides with the introduction of artemisinin combined therapy in Brazil ^15^. Though, *P. falciparum* infections represented on average more than 30% of cases reported during pregnancy, in the assessed years (S2 Table).

**Table 1.**
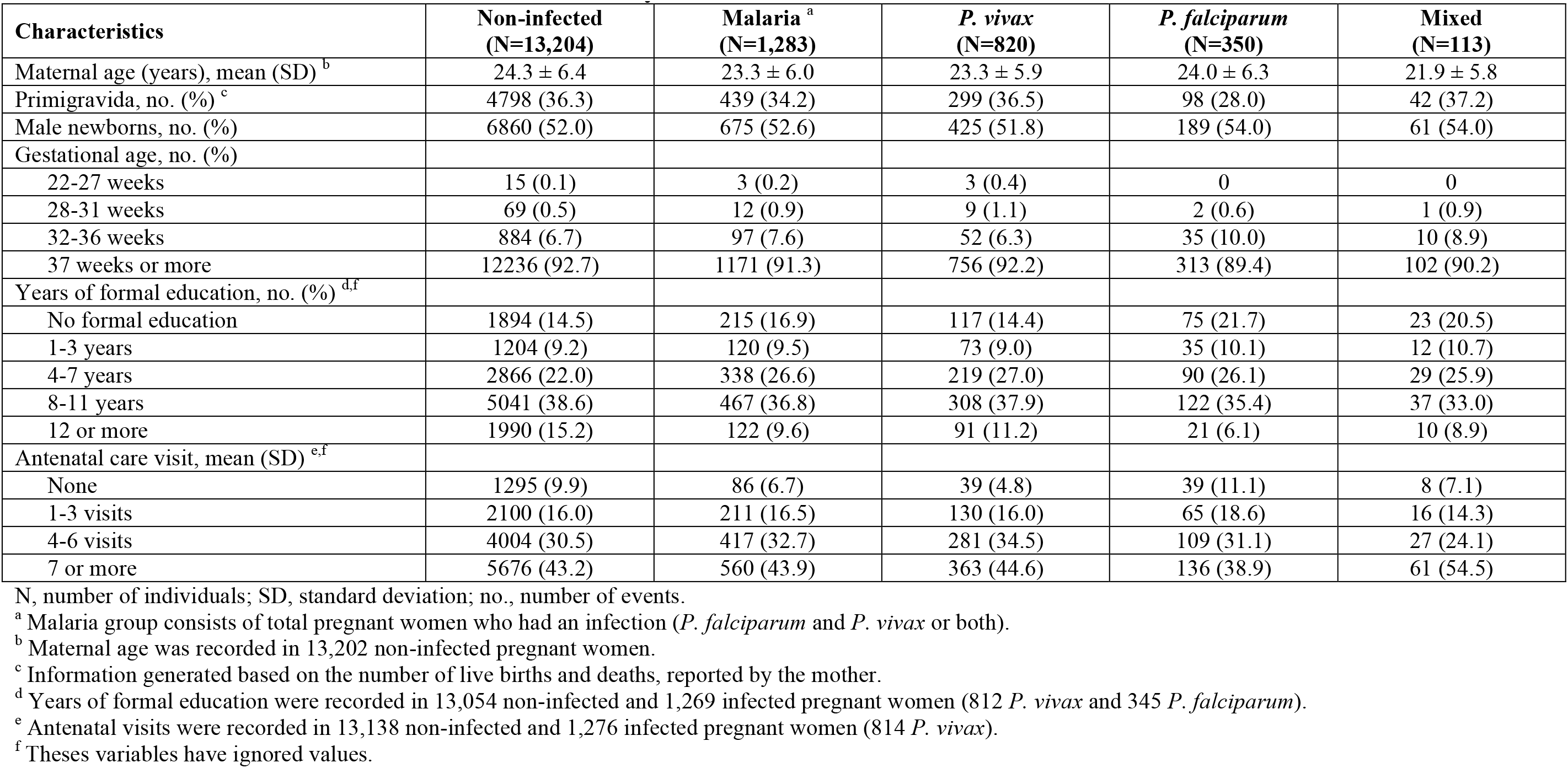
Baseline characteristics of mothers at delivery.

| Characteristics | Non-infected<br>(N=13,204) | Malaria <sup>a</sup><br>(N=1,283) | *P. vivax*<br>(N=820) | *P. falciparum*<br>(N=350) | Mixed<br>(N=113) |
| --- | --- | --- | --- | --- | --- |
| Maternal age (years), mean (SD) <sup>b</sup> | 24.3 ± 6.4 | 23.3 ± 6.0 | 23.3 ± 5.9 | 24.0 ± 6.3 | 21.9 ± 5.8 |
| Primigravida, no. (%) <sup>c</sup> | 4798 (36.3) | 439 (34.2) | 299 (36.5) | 98 (28.0) | 42 (37.2) |
| Male newborns, no. (%) | 6860 (52.0) | 675 (52.6) | 425 (51.8) | 189 (54.0) | 61 (54.0) |
| Gestational age, no. (%) |  |  |  |  |  |
| 22-27 weeks | 15 (0.1) | 3 (0.2) | 3 (0.4) | 0 | 0 |
| 28-31 weeks | 69 (0.5) | 12 (0.9) | 9 (1.1) | 2 (0.6) | 1 (0.9) |
| 32-36 weeks | 884 (6.7) | 97 (7.6) | 52 (6.3) | 35 (10.0) | 10 (8.9) |
| 37 weeks or more | 12236 (92.7) | 1171 (91.3) | 756 (92.2) | 313 (89.4) | 102 (90.2) |
| Years of formal education, no. (%) <sup>d,f</sup> |  |  |  |  |  |
| No formal education | 1894 (14.5) | 215 (16.9) | 117 (14.4) | 75 (21.7) | 23 (20.5) |
| 1-3 years | 1204 (9.2) | 120 (9.5) | 73 (9.0) | 35 (10.1) | 12 (10.7) |
| 4-7 years | 2866 (22.0) | 338 (26.6) | 219 (27.0) | 90 (26.1) | 29 (25.9) |
| 8-11 years | 5041 (38.6) | 467 (36.8) | 308 (37.9) | 122 (35.4) | 37 (33.0) |
| 12 or more | 1990 (15.2) | 122 (9.6) | 91 (11.2) | 21 (6.1) | 10 (8.9) |
| Antenatal care visit, mean (SD) <sup>e,f</sup> |  |  |  |  |  |
| None | 1295 (9.9) | 86 (6.7) | 39 (4.8) | 39 (11.1) | 8 (7.1) |
| 1-3 visits | 2100 (16.0) | 211 (16.5) | 130 (16.0) | 65 (18.6) | 16 (14.3) |
| 4-6 visits | 4004 (30.5) | 417 (32.7) | 281 (34.5) | 109 (31.1) | 27 (24.1) |
| 7 or more | 5676 (43.2) | 560 (43.9) | 363 (44.6) | 136 (38.9) | 61 (54.5) |

N, number of individuals; SD, standard deviation; no., number of events.
<sup>a</sup> Malaria group consists of total pregnant women who had an infection (*P. falciparum* and *P. vivax* or both).
<sup>b</sup> Maternal age was recorded in 13,202 non-infected pregnant women.
<sup>c</sup> Information generated based on the number of live births and deaths, reported by the mother.
<sup>d</sup> Years of formal education were recorded in 13,054 non-infected and 1,269 infected pregnant women (812 *P. vivax* and 345 *P. falciparum*).
<sup>e</sup> Antenatal visits were recorded in 13,138 non-infected and 1,276 infected pregnant women (814 *P. vivax*).
<sup>f</sup> These variables have ignored values.

**Figure 3.**
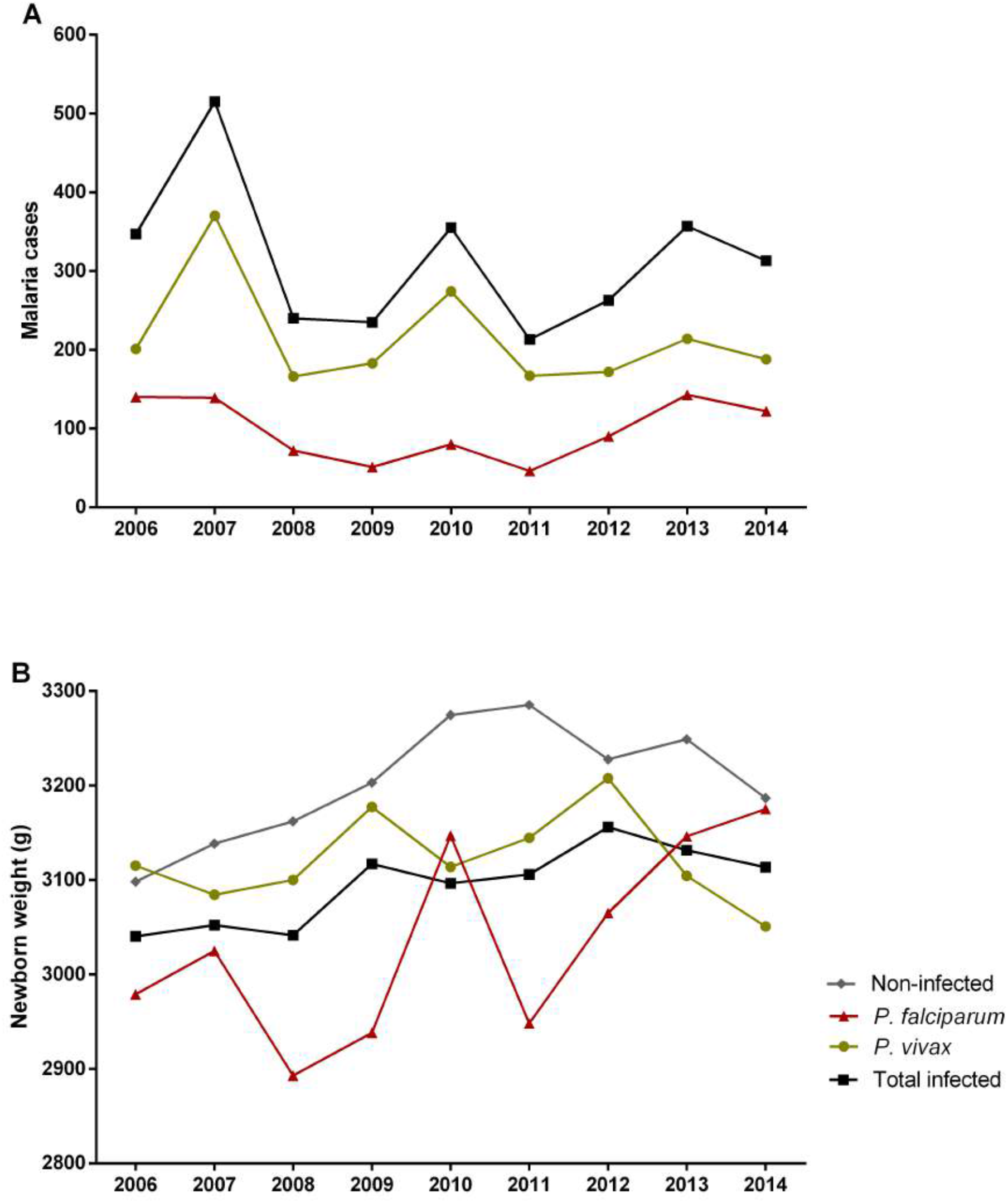
Time-series of gestational malaria cases between 2006-2014. (A) Number of gestational malaria cases per species, (B) mean birth weight of newborns from non-infected and infected women during pregnancy.

### Association of gestational malaria with reduction of the newborns’ birth weight

The analysis of the newborns birth weight across the nine years period, allowed to observe a significant reduction in the mean weight of babies born from women that had malaria during pregnancy (Fig 3B, Table 2, and S3 Table). Newborns from *P. falciparum-infected* mothers presented a more prominent difference of approximately 150 g (p<0.0001) when compared to newborns from non-infected women (Table 2). Notably, the comparison of each group by year evidenced that newborns from *P. vivax-infected* mothers showed higher weight reduction when compared with non-infected (S3 Table). These differences can be explained by the higher prevalence of newborns with LBW among *P. vivax*-infected women (term LBW: NI 4.8%, Pv 6.5%, p=0.031; all LBW: NI 6.8%, Pv 8.9%, p=0.020) (Table 2 and S4 Table). Although this prevalence occurred throughout the assessed years, it was more evident in 2006 and 2013 (S4 Table).

**Table 2.** Clinical outcomes of newborns at birth.

| Characteristics | Non-infected<br>(N=13,204) | Malaria <sup>a</sup><br>(N=1,283) | p value <sup>b</sup> | *P. vivax*<br>(N=820) | p value <sup>b</sup> | *P. falciparum*<br>(N=350) | p value <sup>b</sup> | Mixed<br>(N=113) | p value <sup>b</sup> |
| --- | --- | --- | --- | --- | --- | --- | --- | --- | --- |
| All birth weight (g) |  |  | <0.0001 |  | <0.0001 |  | <0.0001 |  | 0.0002 |
| Mean (SD) | 3200.1 ± 514.7 | 3090.1 ± 524.2 |  | 3118.9 ± 532.4 |  | 3049.9 ± 506.7 |  | 3005.8 ± 503.9 |  |
| Median (IQR) | 3215.0<br>(2900.0-3530.0) | 3100.0<br>(2785.0-3420.0) |  | 3132.5<br>(2800.0-3447.5) |  | 3030.0<br>(2780.0-3350.0) |  | 3095.0<br>(2690.0-3330.0) |  |
| Term birth weight (g) <sup>c</sup> |  |  | <0.0001 |  | <0.0001 |  | <0.0001 |  | 0.004 |
| Mean (SD) | 3236.1 ± 482.2 | 3139.6 ± 471.6 |  | 3159.5 ± 486.0 |  | 3106.4 ± 447.9 |  | 3094.4 ± 426.9 |  |
| Median (IQR) | 3240.0<br>(2940.0-3550.0) | 3130.0<br>(2840.0-3430.0) |  | 3150.0<br>(2842.5-3452.5) |  | 3060.0<br>(2850.0-3380.0) |  | 3155.0<br>(2770.0-3348.0) |  |
| All low birth weight, no. (%) | 896 (6.8) | 120 (9.4) | 0.001 | 73 (8.9) | 0.020 | 31 (8.9) | 0.130 | 16 (14.2) | 0.002 |
| Term low birth weight, no. (%) <sup>c</sup> | 581 (4.8) | 74 (6.3) | 0.017 | 49 (6.5) | 0.031 | 17 (5.4) | 0.575 | 8 (7.8) | 0.144 |
| Prematurity, no. (%) | 968 (7.3) | 112 (8.7) | 0.069 | 64 (7.8) | 0.614 | 37 (10.6) | 0.022 | 11 (9.7) | 0.330 |
| Very preterm birth, no. (%) | 69 (7.1) | 12 (10.7) | 0.058 | 9 (14.1) | 0.032 | 2 (5.4) | 0.901 | 1 (9.1) | 0.596 |
| Late preterm birth, no. (%) | 884 (91.3) | 97 (86.6) | 0.239 | 52 (81.3) | 0.694 | 35 (94.6) | 0.015 | 10 (90.9) | 0.362 |
| Very preterm birth weight (g) |  |  | 0.868 |  | 0.863 |  | 0.627 |  | 0.638 |
| Mean (SD) | 2036.3 ± 722.2 | 2032.3 ± 625.5 |  | 1978.9 ± 692.0 |  | 2112.5 ± 576.3 |  | 2353.0 |  |
| Median (IQR) | 1880.0<br>(1460.0-2575.0) | 2045.0<br>(1532.5-2547.5) |  | 1915.0<br>(1360.0-2575.0) |  | 2112.5<br>(1705.0-2520.0) |  | 2353.0 |  |
| Late preterm birth weight (g) |  |  | 0.093 |  | 0.838 |  | 0.164 |  | 0.0009 |
| Mean (SD) | 2814.1 ± 621.6 | 2683.1 ± 677.2 |  | 2839.0 ± 645.6 |  | 2598.6 ± 703.5 |  | 2167.5 ± 444.1 |  |
| Median (IQR) | 2840.0<br>(2422.5-3232.5) | 2720.0<br>(2240.0-3100.0) |  | 2797.5<br>(2397.5-3267.5) |  | 2765.0<br>(2320.0-3075.0) |  | 2225.0<br>(1860.0-2500.0) |  |

N, number of individuals; no., number of events; SD, standard deviation; IQR, interquartile range.

Further, multivariate logistic regression analysis disclosed the association of malaria with the likelihood of occurring newborns small for gestational age (SGA) at term (weight ≤ 10^th^ centile, boys ≤ 2758g and girls ≤ 2658g) (Odds ratio [OR] 1.23, 95%, confidence interval [CI] 1.05-1.45, p=0.013), which relates to *P. vivax* infection (OR 1.24, 95% CI 1.02-1.52, p=0.035) (Fig 4). Moreover, LBW at term was significantly increased in newborns from malaria-infected mothers, (OR 1.34, 95% CI 1.04-1.72, p=0.024), which was evidenced when mothers were infected by *P. vivax* (OR 1.39, 95% CI 1.03-1.88, p=0.033) (Fig 4). Additionally, segregation by gravidity showed that newborns at term from both primigravida and multigravida presented reduced birth weight when mothers had malaria during pregnancy, irrespective of species (Fig 5A-B, and S5 Table). Nevertheless, newborns from primigravida showed a more prominent birth weight reduction upon infection (Fig 5B, and S5 Table).

**Figure 4.**
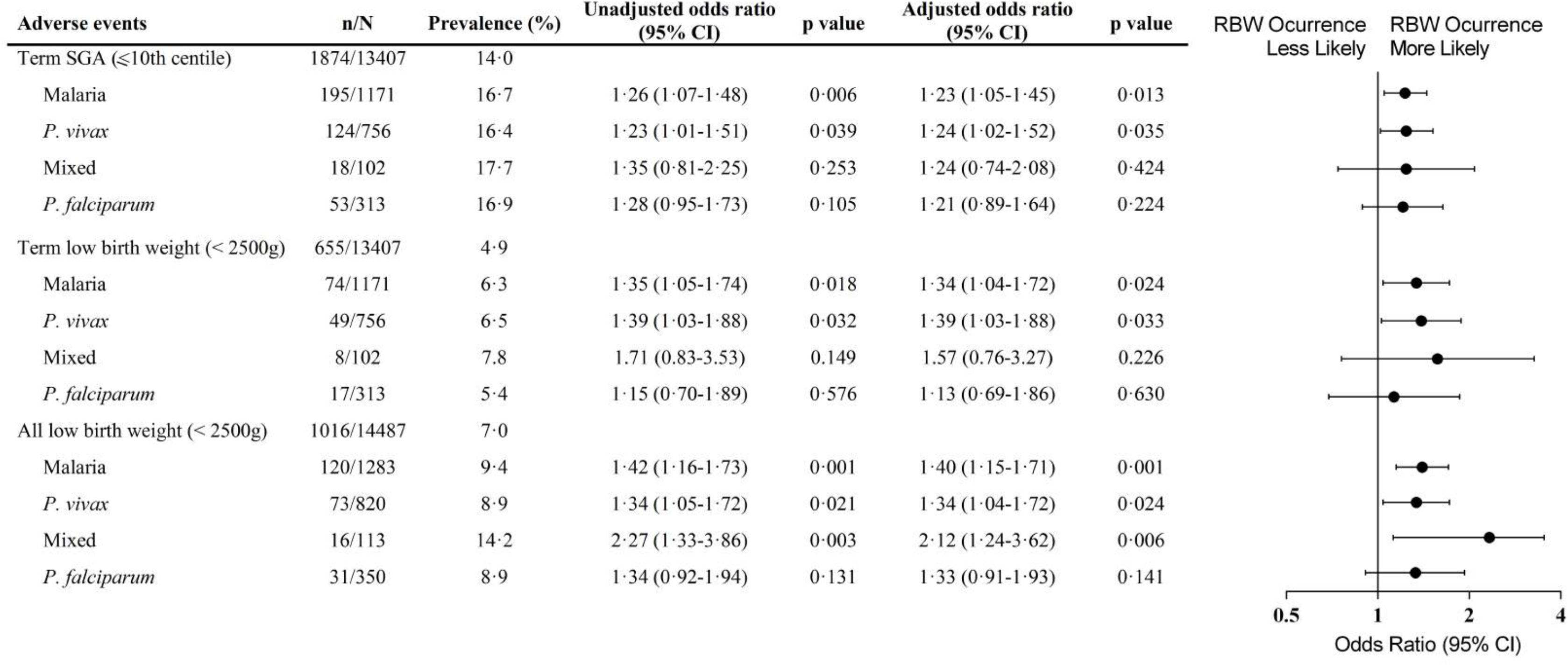
Forest plot of the Odds Ratio for weight reduction in newborns from women infected during pregnancy compared to babies from non-infected women, according to *Plasmodium* species. Each model adjusting for maternal age, parity and years of formal education (less than 4 years); mixed infection *(P. vivax* and *P. falciparum-infection).* p values were estimated through logistic regression methods. n, number of events; N, total number in each group; CI, confidence interval; SGA, small for gestational age; LBW, low birth weight.

**Figure 5.**
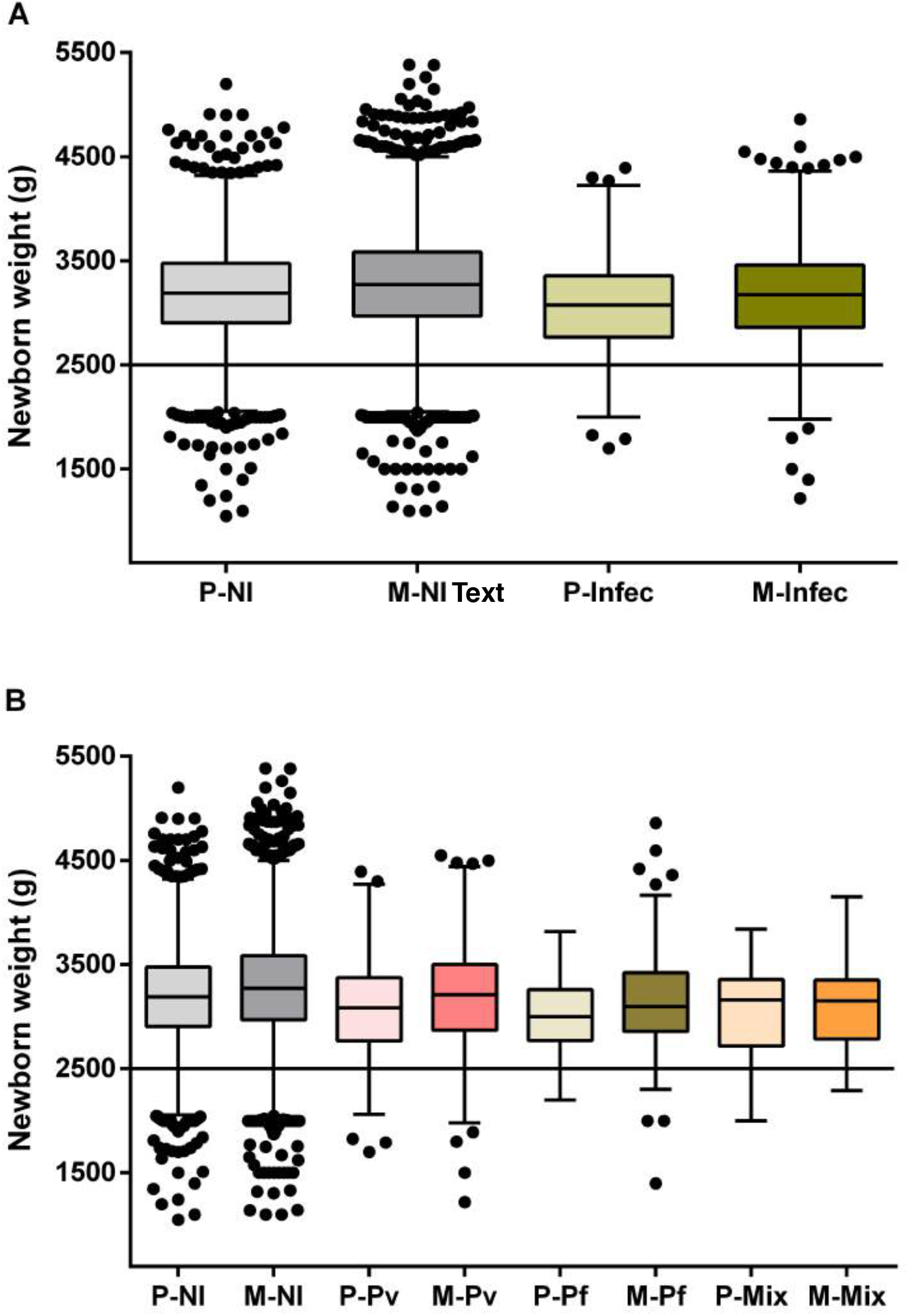
Impact of malaria on birth weight at term according to gravidity. Tukey boxplots show the gravidity effect on the weight of newborns from malaria-infected women (A), and on newborns from women infected according with *Plasmodium* species (B). The bottom and the top of the box are the first and third quartiles, the line inside the box is the median, and the whiskers represent the lowest and the highest data within 1.5 IQR of the first and upper quartiles. The line indicates the cut-off of low birth weight. Differences between each group were examined with Mann-Whitney or Kruskal-Wallis test with a Dunn’s post hoc test. (A) P - NI x Infec (p<0.0001); M - NI x Infec (p<0.0001); NI- P x M (p<0.0001); and Infec - P x M (p=0.0004). (B) P - NI x Pv (p=0.0001); NI x Pf (p=0.0003); M - NI x Pv (p=0.0009); NI x Pf (p<0.0001); NI x Mix (p=0.003); Pv x Pf (p=0.025); Pv - P x M (p=0.0009). P, primigravida; M, multigravida; NI, non-infected pregnant women; Infec, infected pregnant women; Pv, *P. vivax*-infection; Pf, *P. falciparum-* infection; Mix, mixed-infection.

### *P. falciparum* infection during pregnancy increases preterm births

The assembly of databases unveiled increased prematurity among babies born from *P. falciparum-infected* women during pregnancy (Table 2). Prematurity prevalence increased around 3% when women were infected with *P. falciparum*, and the association was evidenced by multivariate logistic regression analysis (OR 1.54, 95% CI 1.09-2.18, p=0.016), which corresponded with late preterm births (OR 1.59, 95% CI 1.11-2.27, p=0.011) (Fig 6). Moreover, *P. vivax* infections were related to very preterm births in women with malaria during pregnancy (OR 2.09, 95% CI 1.04-4.20, p=0.039) (Fig 6).

**Figure 6.**
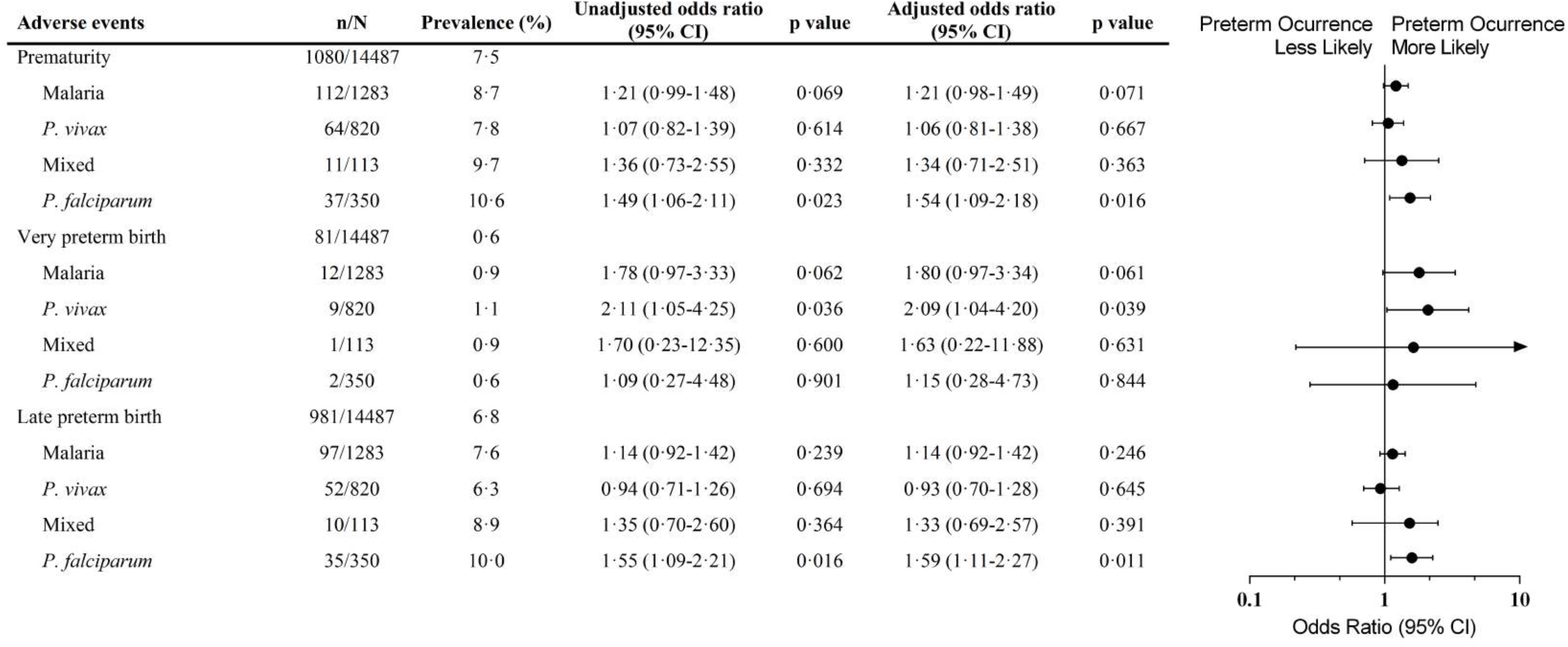
Forest plot of the Odds Ratio for prematurity in newborns from women infected during pregnancy compared to babies from non-infected women, according to *Plasmodium* species. Each model adjusting for maternal age, parity and years of formal education (less than 4 years). Mixed infection (*P. vivax* and *P. falciparum*-infection). p values were estimated through logistic regression methods. n, number of events; N, total number in each group; CI, confidence interval.

Together, these results demonstrate that linkage of national record databases is a valuable research tool, which disclosed adverse neonatal outcomes upon malaria infection during pregnancy in Brazil.

## DISCUSSION

Malaria during pregnancy is known as an important risk factor for miscarriage, stillbirth, LBW and maternal anemia ^8,21-23^. Nevertheless, little is known about malaria in pregnancy in the Americas endemic regions, where it predominates *P. vivax* infections. This work is the first to assess the effect of gestational malaria in Brazil through the linkage of national databases of the Brazilian Ministry of Health, SINASC and SIVEP-Malaria. Interestingly, despite the decrease in the number of gestational malaria cases through the studied period (2006-2014), the impact of malaria during pregnancy is still evident. The reduction of the mean of the weight was maintained throughout the years and the higher prevalence of preterm births among newborns from women that presented malaria during pregnancy.

The number of studies estimating the real frequency of malaria in pregnant women is still limited, both in Brazil and in other regions of the Americas, which are considered low transmission areas ^2^. The prevalence (8.9%) of malaria during pregnancy in our study was similar to the findings of a multi-centric study that enrolled data from the Americas (Guatemala, Colombia, and Brazil) between 2008-2011, and another study performed in Urubá (Colombia) between 2005-2009 ^24,25^. Though, the prevalence is higher in relation to reports from Iquitos (6.6%) (Peru), and from other Brazilian cities, such as Manaus (6.1%) and Coari (4.3%) in the Amazonas state, and Rio Branco (1.4%) in Acre state ^26-29^. The discrepancies may encompass differential study designs and endemicity of studied areas.

Prematurity is one of the adverse effects commonly observed in malaria during pregnancy ^6,30,31^. Usually, it correlates with infections occurring during the third trimester of pregnancy and contributes to increasing the number of newborns with LBW, which is more likely to be observed in low transmission areas ^2,32^. In fact, our data show that *P. falciparum* infections during pregnancy are responsible for a high proportion of preterm births, mainly late preterm births (≥32 and <37 weeks of gestation). However, it was not possible to correlate the time of infection with the gestational trimester.

Newborns reduced weight at birth either classified as LBW or SGA, is an important predictive marker of neonatal and child survival, and can result from two basic factors: intrauterine growth restriction and preterm births ^33,31^. In gestational malaria, birth weight reduction is the main adverse outcome observed in studies involving *P. falciparum* infections^2,30,34,35^. In our observations, malaria infection during pregnancy represents a critical morbidity that impacts newborns’ weight. The records show that malaria in pregnant women increases the number of babies born at term with SGA by 19.3% and LBW by 28.6% (SGA - NI 14.0%, Malaria 16.7%; term LBW - NI 4.9%, Malaria 6.3%). Strikingly, *P. vivax* infection during pregnancy represents the higher odds for the occurrence of birth weight reduction (SGA - OR 1.24, 95% CI 1.02-1.52, p=0.035; term LBW - OR 1.39, 95% CI 1.03-1.88, p=0.033). The absence of the association between *P. falciparum* infections and the occurrence of SGA or LBW newborns can be related with the restricted pool of variant genes of the Amazonian parasite, which can explain the mild outcomes observed in the Americas, substantially different from other endemic regions in the world ^36^. Nevertheless, our data corroborate some findings in Southeast Asia from Moore *et al.* that show that *P. vivax* infection is associated with SGA and *P. falciparum* infection with late preterm, although we could not correlate with time of infection ^34^.

The reduction of newborns birth weight is multifactorial, and it can be related to social-economic, environmental, nutritional, and clinic factors during pregnancy. However, in this study, it was not possible to assess other risk factors, once these variables were absent in the databases used. Of note, it is important to highlight that it is impossible to compare this study with other carried out in Africa. There, *P. falciparum* infections are predominant and, in general, the health systems that diagnose and treat malaria have several limitations, summed up with the high rate of co-infection with other diseases, such as HIV and tuberculosis. The Brazilian Amazonian region has a health care system with effective strategies to control, diagnose, and treat malaria, despite being a low transmission area with predominance of *P. vivax* infections. These characteristics make our findings even more interesting, as we observed a substantial impact of infection during pregnancy in newborns.

In Brazil, malaria is a mandatory notification disease, and SIVEP-Malaria is essential to plan health interventions that enable effective control and preventive strategies to eradicate the disease. For pregnant women, the early diagnosis is essential to prevent adverse outcomes. In 2014, it was enforced, by Brazilian MoH, a malaria routine screen during antenatal care and at delivery, in women living in Brazilian Amazonian region states ^37^. This initiative brought important benefits for both mother and fetus, enabling early treatment and preventing gestational adverse outcomes.

This work present potential limitations. First, the databases used have only two shared variables with adequate fulfillment, and this hampered the identification of all women. Therefore, the number of pregnant women with malaria can be underestimated. Second, the reduction of birth weight has different etiologies. Although we used important exclusion criteria, it was not possible to identify through SINASC women presenting other infections, such as TORCHs, as well as, nutritional or other risk factors.

In conclusion, this work allowed us to observe through a time-lapse study, the effect of gestational malaria on newborns birth weight in a region considered of low transmission and with *P. vivax* infections predominance. During the evaluated period (2006-2014), malaria infections continue to be an important risk factor for prematurity and reduction of newborns’ birth weight, despite the decline in the number of cases reported in the region. We have shown that the SINASC and the SIVEP-Malaria databases linkage allow to estimate the extent of malaria adverse effects, which permit to improve information and further plan interventions. These findings reinforce the urgent need for health programs and actions to prevent and protect pregnant women against the consequences of malaria, especially during the antenatal care.

## Acknowledgement

We thank Health Surveillance Secretariat of Acre for authorizing the data collection. Also, we thank the Municipal Health Secretariat of Cruzeiro do Sul, which promptly welcome us and provided us access to the SINASC database, and to the Brazilian Epidemiological Surveillance/Administration of Endemics, which authorized the assess to SIVEP-Malaria information.

## Contributors

JGD, LAG, and CRFM designed the study and were involved in data acquisition and scientific input. JGD, RMS, NRMS, AB, SE, LAG, and CRFM contributed to the analysis and interpretation of data. JGD, LAG, and CRFM wrote the manuscript. CRFM and SE were the main funders of this work. CRFM had full access to all the data in the study and takes responsibility for the integrity of the data and the accuracy of the data analysis. All authors reviewed and approved the final version of this manuscript.

## SUPPORTING INFORMATION CAPTIONS

**S1 Table.** Characteristics of mothers and newborns per year in Cruzeiro do Sul, 2006-2014. Data from the System Information of Live Births provided by the Cruzeiro do Sul Municipal Secretariat of Health. ^a^ Information generated based on the number of live births and deaths reported by the mother. ^b^ There are other groups with ignored values.

| Characteristics | 2006<br>(N=1594) | 2007<br>(N=1795) | 2008<br>(N=1748) | 2009<br>(N=1625) | 2010<br>(N=1382) | 2011<br>(N=1510) | 2012<br>(N=1553) | 2013<br>(N=1620) | 2014<br>(N=1660) |
| --- | --- | --- | --- | --- | --- | --- | --- | --- | --- |
| **Age, years, mean (SD)** | 23.6 (6.1) | 23.7 (6.1) | 24.4 (6.3) | 24.5 (6.3) | 24.3 (6.3) | 24.3 (6.5) | 24.1 (6.4) | 24.3 (6.4) | 24.4 (6.6) |
| **Primigravida, no. (%)<sup>a</sup>** | 551 (34.6) | 481 (26.8) | 534 (30.6) | 530 (32.6) | 520 (37.6) | 644 (42.7) | 632 (40.7) | 659 (40.7) | 688 (41.5) |
| **Gestational age, no. (%)** |  |  |  |  |  |  |  |  |  |
| 22-27 weeks | 1 (0.1) | 5 (0.3) | 3 (0.2) | - | - | 5 (0.3) | - | 1 (0.1) | 3 (0.2) |
| 28-31 weeks | 5 (0.3) | 2 (0.1) | 7 (0.4) | 2 (0.1) | 11 (0.8) | 22 (1.5) | 16 (1.0) | 9 (0.5) | 7 (0.4) |
| 32-36 weeks | 27 (1.7) | 34 (1.9) | 55 (3.1) | 68 (4.2) | 64 (4.6) | 157 (10.4) | 199 (12.8) | 164 (10.1) | 213 (12.8) |
| 37 weeks or more | 1561 (97.9) | 1754 (97.7) | 1683 (96.3) | 1555 (95.7) | 1307 (94.6) | 1326 (87.8) | 1338 (86.2) | 1446 (89.3) | 1437 (86.6) |
| **Marital status, no. (%)<sup>b</sup>** |  |  |  |  |  |  |  |  |  |
| Single | 1408 (89.1) | 1503 (84.3) | 1390 (81.0) | 1329 (83.3) | 1021 (75.3) | 516 (34.8) | 414 (26.8) | 378 (23.6) | 369 (22.3) |
| Married or cohabiting | 163 (10.3) | 257 (14.4) | 310 (18.1) | 252 (15.8) | 327 (24.1) | 960 (64.6) | 1120 (72.6) | 1199 (74.9) | 1272 (76.8) |
| **Years of formal education, no. (%)<sup>b</sup>** |  |  |  |  |  |  |  |  |  |
| No education | 616 (38.8) | 622 (34.9) | 248 (14.4) | 202 (12.7) | 130 (9.5) | 86 (5.8) | 66 (4.3) | 77 (4.8) | 62 (3.8) |
| 1-3 years | 120 (7.6) | 135 (7.6) | 282 (16.4) | 239 (15.0) | 170 (12.5) | 168 (11.2) | 92 (6.0) | 63 (3.9) | 55 (3.4) |
| 4-7 years | 226 (14.2) | 264 (14.8) | 487 (28.3) | 469 (29.4) | 383 (28.1) | 390 (26.1) | 371 (24.0) | 317 (19.8) | 297 (18.1) |
| 8-11 years | 402 (25.3) | 295 (16.6) | 483 (28.1) | 477 (29.9) | 461 (33.9) | 675 (45.2) | 815 (52.7) | 914 (57.1) | 986 (60.2) |
| 12 or more | 222 (14.0) | 461 (25.9) | 219 (12.7) | 206 (12.9) | 217 (15.9) | 170 (11.4) | 196 (12.7) | 206 (12.9) | 215 (13.1) |
| **Antenatal visits, no. (%)<sup>b</sup>** |  |  |  |  |  |  |  |  |  |
| None | 245 (15.6) | 233 (13.1) | 295 (16.9) | 241 (15.0) | 118 (8.6) | 101 (6.7) | 57 (3.7) | 57 (3.5) | 34 (2.0) |
| 1-3 visits | 147 (9.4) | 132 (7.4) | 332 (19.1) | 399 (24.8) | 366 (26.7) | 285 (18.9) | 235 (15.1) | 213 (13.2) | 202 (12.2) |
| 4-6 visits | 216 (13.7) | 191 (10.8) | 583 (33.5) | 568 (35.4) | 552 (40.2) | 621 (41.1) | 591 (38.1) | 556 (34.3) | 543 (32.7) |
| 7 or more | 960 (61.1) | 1213 (68.3) | 523 (30.0) | 380 (23.7) | 314 (22.9) | 502 (33.3) | 669 (43.1) | 794 (49.0) | 881 (53.1) |
| **Caesarean section, no. (%)** | 317 (19.9) | 418 (23.3) | 469 (26.9) | 509 (31.3) | 543 (39.3) | 525 (34.8) | 623 (40.1) | 744 (45.9) | 752 (45.3) |
| **Birth weight (g), mean (SD)** |  |  |  |  |  |  |  |  |  |
| Mean (SD) | 3093.3 (523.6) | 3128.3 (529.0) | 3150.8 (506.7) | 3197.3 (509.6) | 3256.2 (524.0) | 3271.7 (503.7) | 3222.7 (515.0) | 3237.5 (486.2) | 3180.8 (521.9) |
| Median (IQR) | 3100.0 (2700.0-3450.0) | 3450.0 (2770.0-3480.0) | 3170.0 (2850.0-3500.0) | 3220.0 (2900.0-3515.0) | 3270.0 (2950.0-3590.0) | 3275.0 (2990.0-3585.0) | 3225.0 (2930.0-3560.0) | 3235.0 (2955.0-3537.5) | 3210.0 (2907.5-3500.0) |
| **Low birth weight, no. (%)** | 123 (7.7) | 134 (7.5) | 126 (7.2) | 111 (6.8) | 96 (7.0) | 80 (5.3) | 114 (7.3) | 95 (5.9) | 137 (8.3) |
| **Very low birth weight, no. (%)** | 6 (0.4) | 11 (0.6) | 10 (0.6) | 4 (0.3) | 4 (0.3) | 5 (0.3) | 7 (0.5) | 7 (0.4) | 17 (1.0) |

**S2 Table.** Trends in malaria infection during pregnancy in Cruzeiro do Sul, 2006-2014. N, number of malaria cases. Data are N or N (%). Values correspond to the total number of malaria episodes reported between 2006 and 2014.

| Cases of malaria | 2006 | 2007 | 2008 | 2009 | 2010 | 2011 | 2012 | 2013 | 2014 |
| --- | --- | --- | --- | --- | --- | --- | --- | --- | --- |
| Overall | 347 | 515 | 240 | 235 | 355 | 213 | 263 | 357 | 313 |
| *P. falciparum* | 140 (40.4%) | 139 (27.0%) | 72 (30.0%) | 51 (21.7%) | 80 (22.5%) | 46 (21.6%) | 90 (34.2%) | 143 (40.1%) | 122 (39.0%) |
| *P. vivax* | 201 (57.9%) | 370 (71.8%) | 166 (69.2%) | 183 (77.9%) | 274 (77.2%) | 167 (78.4%) | 172 (65.4%) | 214 (59.9%) | 188 (60.0%) |
| Mixed | 6 (1.7%) | 6 (1.2%) | 2 (0.8%) | 1 (0.4%) | 1 (0.3%) | 0 | 1 (0.4%) | 0 | 3 (1.0%) |

**S3 Table.** Description of the birth weight of newborns from Non-Infected and Infected pregnant women per year. N, number of individuals; IQR, interquartile range. ^a^ Malaria group consists of total pregnant women who had an infection (*P. falciparum*, *P. vivax*, and Mixed infections). ^b^ Differences between Non-Infected and the other groups were evaluated using Mann-Whitney rank sum tests.

| Year | Non-infected |  | Malaria <sup>a</sup> |  | p value<sup>b</sup> | *P. vivax* |  | p value<sup>b</sup> | *P. falciparum* |  | p value<sup>b</sup> | Mixed |  | p value<sup>b</sup> |
| --- | --- | --- | --- | --- | --- | --- | --- | --- | --- | --- | --- | --- | --- | --- |
|  | N | Median (IQR) | N | Median (IQR) |  | N | Median (IQR) |  | N | Median (IQR) |  | N | Median (IQR) |  |
| 2006 | 1459 | 3100.0 (2700.0-3450.0) | 135 | 3030.0 (2660.0-3370.0) | 0.174 | 72 | 3135.0 (2770.0-3455.0) | 0.071 | 51 | 3005.0 (2615.0-3300.0) | 0.623 | 12 | 2827.5 (2612.5-3350.0) | 0.259 |
| 2007 | 1581 | 3170.0 (2795.0-3500.0) | 214 | 3077.5 (2690.0-3400.0) | 0.047 | 147 | 3100.0 (2720.0-3445.0) | 0.245 | 38 | 3017.5 (2640.0-3300.0) | 0.816 | 29 | 3060.0 (2500.0-3340.0) | 0.087 |
| 2008 | 1586 | 3190.0 (2860.0-3500.0) | 162 | 3000.0 (2730.0-3390.0) | 0.270 | 99 | 3100.0 (2770.0-3405.0) | 0.521 | 45 | 2990.0 (2600.0-3165.0) | 0.482 | 18 | 3055.0 (2770.0-3300.0) | 0.338 |
| 2009 | 1517 | 3230.0 (2910.0-3530.0) | 108 | 3117.5 (2850.0-3417.5) | 0.559 | 83 | 3210.0 (2880.0-3460.0) | 0.826 | 22 | 2900.0 (2750.0-3000.0) | 0.664 | 3 | 2660.0 (2415.0-3200.0) | 0.102 |
| 2010 | 1239 | 3300.0 (2970.0-3610.0) | 143 | 3090.0 (2785.0-3385.0) | 0.108 | 110 | 3095.0(2790.0-3430.0) | 0.033 | 24 | 3185.0 (2995.0-3342.5) | 0.362 | 9 | 2630.0 (2420.0-2920.0) | 0.002 |
| 2011 | 1394 | 3287.5 (3010.0-3600.0) | 116 | 3122.5 (2780.0-3420.0) | 0.009 | 93 | 3150.0 (2800.0-3425.0) | 0.008 | 18 | 2972.5 (2670.0-3245.0) | 0.472 | 5 | 2880.0 (2760.0-3265.0) | 0.133 |
| 2012 | 1442 | 3232.5 (2930.0-3560.0) | 111 | 3100.0 (2845.0-3475.0) | 0.107 | 73 | 3200.0 (2870.0-3480.0) | 0.629 | 30 | 3030.0 (2745.0-3340.0) | 0.463 | 8 | 3075.0 (2865.0-3190.0) | 0.198 |
| 2013 | 1461 | 3245.0 (2975.0-3540.0) | 159 | 3145.0 (2860.0-3520.0) | 0.091 | 71 | 3210.0 (2810.0-3575.0) | 0.181 | 74 | 3127.5 (2890.0-3475.0) | 0.429 | 14 | 3270.0 (2920.0-3485.0) | 0.772 |
| 2014 | 1525 | 3215.0 (2915.0-3510.0) | 135 | 3160.0 (2855.0-3400.0) | 0.087 | 72 | 3062.5 (2805.0-3395.0) | 0.067 | 48 | 3205.0 (2957.5-3410.0) | 0.932 | 48 | 3300.0 (3140.0-3470.0) | 0.715 |

**S4 Table.**
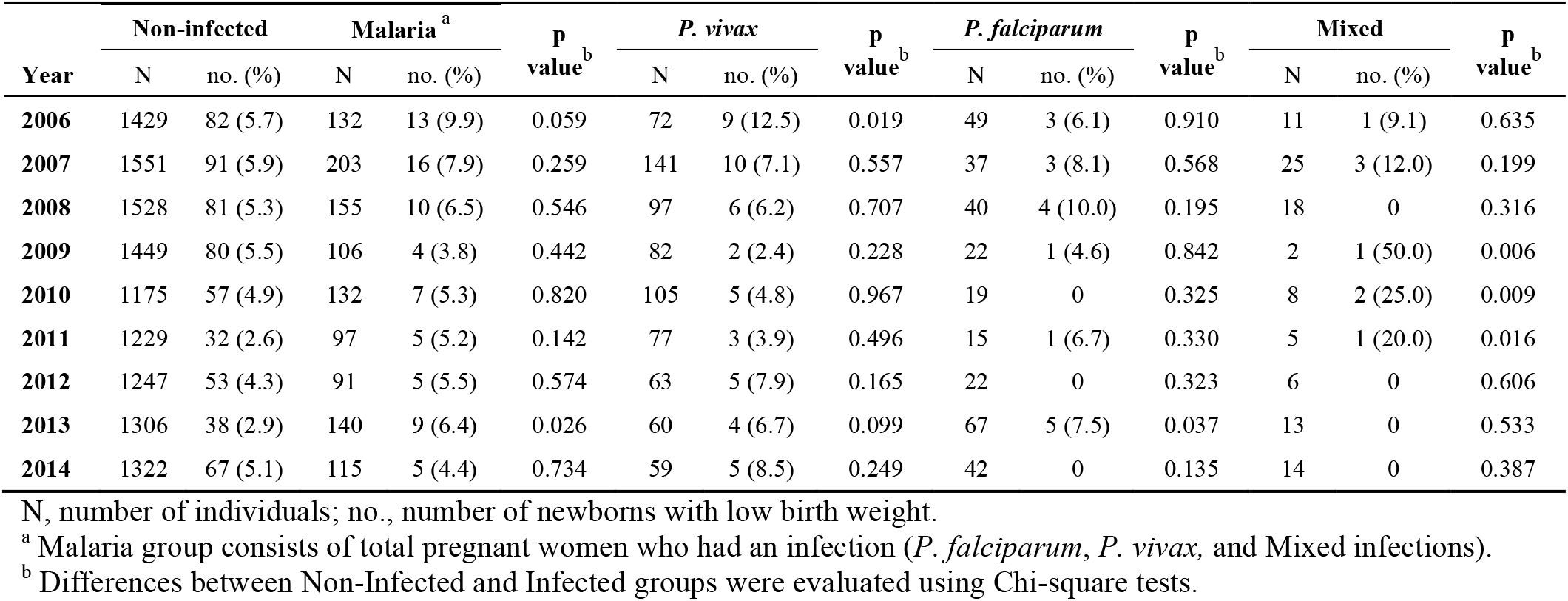
Description of term low birth weight newborns from Non-Infected and Infected pregnant women per year. N, number of individuals; no., number of newborns with low birth weight. ^a^ Malaria group consists of total pregnant women who had an infection (*P. falciparum*, *P. vivax*, and Mixed infections). ^b^ Differences between Non-Infected and Infected groups were evaluated using Chi-square tests.

| Year | Non-infected |  | Malaria<sup>a</sup> |  | p value<sup>b</sup> | *P. vivax* |  | p value<sup>b</sup> | *P. falciparum* |  | p value<sup>b</sup> | Mixed |  | p value<sup>b</sup> |
| --- | --- | --- | --- | --- | --- | --- | --- | --- | --- | --- | --- | --- | --- | --- |
|  | N | no. (%) | N | no. (%) |  | N | no. (%) |  | N | no. (%) |  | N | no. (%) |  |
| 2006 | 1429 | 82 (5.7) | 132 | 13 (9.9) | 0.059 | 72 | 9 (12.5) | 0.019 | 49 | 3 (6.1) | 0.910 | 11 | 1 (9.1) | 0.635 |
| 2007 | 1551 | 91 (5.9) | 203 | 16 (7.9) | 0.259 | 141 | 10 (7.1) | 0.557 | 37 | 3 (8.1) | 0.568 | 25 | 3 (12.0) | 0.199 |
| 2008 | 1528 | 81 (5.3) | 155 | 10 (6.5) | 0.546 | 97 | 6 (6.2) | 0.707 | 40 | 4 (10.0) | 0.195 | 18 | 0 | 0.316 |
| 2009 | 1449 | 80 (5.5) | 106 | 4 (3.8) | 0.442 | 82 | 2 (2.4) | 0.228 | 22 | 1 (4.6) | 0.842 | 2 | 1 (50.0) | 0.006 |
| 2010 | 1175 | 57 (4.9) | 132 | 7 (5.3) | 0.820 | 105 | 5 (4.8) | 0.967 | 19 | 0 | 0.325 | 8 | 2 (25.0) | 0.009 |
| 2011 | 1229 | 32 (2.6) | 97 | 5 (5.2) | 0.142 | 77 | 3 (3.9) | 0.496 | 15 | 1 (6.7) | 0.330 | 5 | 1 (20.0) | 0.016 |
| 2012 | 1247 | 53 (4.3) | 91 | 5 (5.5) | 0.574 | 63 | 5 (7.9) | 0.165 | 22 | 0 | 0.323 | 6 | 0 | 0.606 |
| 2013 | 1306 | 38 (2.9) | 140 | 9 (6.4) | 0.026 | 60 | 4 (6.7) | 0.099 | 67 | 5 (7.5) | 0.037 | 13 | 0 | 0.533 |
| 2014 | 1322 | 67 (5.1) | 115 | 5 (4.4) | 0.734 | 59 | 5 (8.5) | 0.249 | 42 | 0 | 0.135 | 14 | 0 | 0.387 |

**S5 Table.** Description of birth weight of term newborns from Non-Infected and Infected pregnant women per gravidity. N, number of individuals; SD, standard deviation; IQR, interquartile range. ^a^ Malaria group consists of total pregnant women who had an infection (*P. falciparum*, *P. vivax*, and Mixed infections). ^b^ Differences between each group were examined using Mann-Whitney or Kruskal-Wallis test with Dunn post hoc test. ^c^ p=0.025 for *P. vivax* versus *P. falciparum* groups.

| Birth weight (g) | Non-infected<br>(N=12,236) | Malaria <sup>a</sup><br>(N=1,171) | p value <sup>b</sup> | *P. vivax*<br>(N=756) | p value <sup>b</sup> | *P. falciparum*<br>(N=313) | p value <sup>b</sup> | Mixed<br>(N=102) | p value <sup>b</sup> |
| --- | --- | --- | --- | --- | --- | --- | --- | --- | --- |
| Primigravida |  |  |  |  |  |  |  |  |  |
| Mean (SD) | 3184.1 (454.8) | 3072.1 (453.1) | <0.0001 | 3088.0 (478.0) <sup>c</sup> | 0.0001 | 3021.6 (370.9) | 0.0003 | 3070.1 (436.6) | 0.104 |
| Median (IQR) | 3186.5 (2905.0-3475.0) | 3075.0 (2769.0-3355.0) |  | 3085.0 (2770.0-3370.0) |  | 3000.0 (2769.0-3260.0) |  | 3160.0 (2720.0-3348.0) |  |
| Multigravida |  |  |  |  |  |  |  |  |  |
| Mean (SD) | 3265.1 (494.5) | 3173.2 (477.2) | <0.0001 | 3199.0 (486.4) <sup>c</sup> | 0.0009 | 3137.0 (469.5) | <0.0001 | 3108.2 (424.1) | 0.003 |
| Median (IQR) | 3270.0 (2970.0-3585.0) | 3175.0 (2860.0-3460.0) |  | 3210.0 (2870.0-3500.0) |  | 3095.0 (2860.0-3420.0) |  | 3150.0 (2800.0-3340.0) |  |
| p value (Primigravida x Multigravida) | <0.0001 | 0.0004 |  | 0.0009 |  | 0.083 |  | 0.862 |  |

